# MRI-based classifier to identify close-to-onset cases in *C9orf72* genetic frontotemporal dementia

**DOI:** 10.1101/2025.10.18.683237

**Authors:** Mahdie Soltaninejad, Yasser Iturria-Medina, Reza Rajabli, Gleb Bezgin, Niki Hosseini-Kamkar, Arabella Bouzigues, Lucy L. Russell, Phoebe H. Foster, Eve Ferry-Bolder, John C. van Swieten, Lize C. Jiskoot, Harro Seelaar, Raquel Sanchez-Valle, Robert Laforce, Caroline Graff, Daniela Galimberti, Rik Vandenberghe, Alexandre de Mendonça, Pietro Tiraboschi, Isabel Santana, Alexander Gerhard, Johannes Levin, Benedetta Nacmias, Markus Otto, Maxime Bertoux, Thibaud Lebouvier, Chris R. Butler, Isabelle Le Ber, Elizabeth Finger, Maria Carmela Tartaglia, Mario Masellis, James B. Rowe, Matthis Synofzik, Fermin Moreno, Barbara Borroni, Jonathan D. Rohrer, Simon Ducharme, GENFI

**Author notes:** The list of GENFI consortium members is provided in Appendix.

## Abstract

Predicting symptom onset in genetic frontotemporal dementia (FTD) is crucial for advancing targeted interventions and clinical trial design. Brain changes begin years before clinical symptoms emerge, making neuroimaging a strong candidate for onset prediction. However, FTD is highly heterogeneous, encompassing diverse molecular pathologies, affected brain networks, and symptom trajectories. This variability limits the predictive power of any single imaging biomarker and underscores the need for an integrative, multimodal approach to improve prediction accuracy and generalizability.

We used machine learning to integrate diverse neuroimaging features, identifying a robust signature for risk stratification. We analyzed T1-weighted and T2-weighted MRI scans from 71 symptomatic *C9orf72* carriers, 90 presymptomatic carriers, and 69 healthy controls from the GENFI cohort. We used FreeSurfer to measure cortical thickness and subcortical volumes, and BISON to quantify white matter hyperintensities (WMH). We applied Principal Component Analysis for dimensionality reduction and trained a random forest classifier to distinguish symptomatic carriers from controls. The model was subsequently applied to the presymptomatic cohort to identify individuals whose brain patterns resembled those of symptomatic cases, under the hypothesis that greater similarity indicated a higher risk of conversion. We validated the model with neuropsychological data and a two-year longitudinal follow-up.

The classifier distinguished symptomatic *C9orf72* carriers from controls with 87.0% accuracy. When applied to presymptomatic carriers, the model identified 21.1% of the cohort as having brain features comparable to those of symptomatic cases. This “high-risk group” showed significant neuropsychological weaknesses in executive function, language and social cognition compared to the non high-risk group. The model accurately predicted clinical conversion within a two-year period with 84.5% accuracy, a 70% sensitivity and a 93.3% negative predictive value.

Our findings demonstrate the utility of a machine learning approach using multi-modal MRI to identify presymptomatic *C9orf72* carriers at high risk of disease onset within the next two years. By capturing subtle neuroanatomical patterns associated with disease processes, this approach offers a promising method for stratifying genetic FTD carriers prior to symptom onset. Such predictive models could optimize patient selection in future clinical trials.

## 1 Introduction

Approximately one-third of frontotemporal dementia (FTD) cases are inherited, most commonly due to autosomal dominant mutations in *C9orf72*, *GRN*, and *MAPT*.^1^ Among these, *C9orf72* repeat expansion is the most frequent cause of genetic FTD.^2^ Carriers of pathogenic expansions face a near-certain risk of developing disease, yet the age of onset is highly variable, ranging from the 30s to the 80s.^3,4^ The period during which a person carries a pathogenic mutation but has not yet developed clinical symptoms is referred to as the presymptomatic phase. This interval lasts decades, but typically ends with the eventual onset of FTD or amyotrophic lateral sclerosis (ALS) in many carriers. Identifying individuals nearing the transition to the symptomatic stage is therefore of critical importance for both patient care and therapeutic development,^5^ as it would allow families to plan and enable timely recruitment for clinical trials.

Neuroimaging has emerged as a particularly promising source of biomarkers in genetic FTD. Multiple studies demonstrate that structural brain alterations can precede clinical onset by 5–10 years.^6^ In presymptomatic *C9orf72* carriers, structural MRI studies have reported gray matter loss in the insula, thalamus, frontal, temporal, parietal, and occipital cortices, along with subcortical and cerebellar involvement.^6–12^ Several studies highlighted early thalamic atrophy, particularly in the pulvinar and lateral geniculate nuclei, as well as reductions in basal ganglia and medial temporal regions including the amygdala and hypothalamus.^10–14^ White matter abnormalities have also been consistently described, with reduced integrity in corticospinal, thalamic, and frontotemporal tracts, early alterations detectable with advanced diffusion metrics,^8,15–17^ and increased white matter hyperintensity burden in the frontal and occipital lobes.^18^ Finally, ventricular expansion has been observed up to four years before expected symptom onset,^19^ further supporting the presence of widespread structural alterations in *C9orf72* carriers years before clinical conversion. These convergent results suggest that neuroimaging biomarkers can sensitively capture early pathological processes and thus hold strong potential for prognostic applications.

Nevertheless, the heterogeneous clinical and neuroanatomical manifestations of FTD^1,2^ complicate attempts to predict disease progression using single biomarkers alone. Conventional univariate approaches are often insufficient to disentangle the complex, multidimensional changes underlying disease onset. Machine learning (ML) methods offer a promising analytical framework, as they can integrate diverse neuroimaging features into predictive models, handle high-dimensional data, and detect latent disease patterns not apparent in traditional analyses.^20^ Such models are particularly well suited to identifying the subtle “disease signature” that may characterize individuals approaching symptom onset.

In the present study, we sought to develop an MRI-based classifier to identify presymptomatic *C9orf72* carriers at high risk of imminent conversion. We first trained a model to discriminate symptomatic *C9orf72* cases from healthy controls, thereby extracting a robust imaging-based signature of disease. We then applied this classifier to presymptomatic mutation carriers and evaluated whether those assigned a “disease-like” profile were more likely to develop clinical symptoms within two years. Finally, to enhance interpretability, we performed feature-importance analysis to identify which imaging features most strongly influenced model predictions. By leveraging ML and multimodal neuroimaging, this study offers new insight into early detection strategies for genetic FTD and contributes to the design of future preventive interventions.

## 2 Method

### 2.1 Study Design and Participants

This study was conducted using data from the Genetic Frontotemporal Dementia Initiative (GENFI).^6^ The study specifically targeted individuals with the *C9orf72* gene mutation, which is the largest group in GENFI. The cohort included three participant groups: symptomatic carriers, who met established clinical criteria for one of the various FTD syndromes; presymptomatic carriers who carried the mutation but had not yet developed symptoms; and non-carrier controls who are first-degree relatives without the mutation who served as a baseline comparison group.

Participants were included based on the following criteria: a minimum age of 40 years, and the availability of MRI scans and demographic data. The final study population consisted of 235 participants, comprising 80 symptomatic carriers, 82 presymptomatic carriers, and 73 non-carrier controls. Detailed inclusion and exclusion numbers are presented in Figure 1. The study was approved by the Research Ethics Board of the McGill University Health Centre (MP-20-2016-2500) and by local ethics committees at participating sites, with written informed consent obtained from all participants.

**Figure 1.**
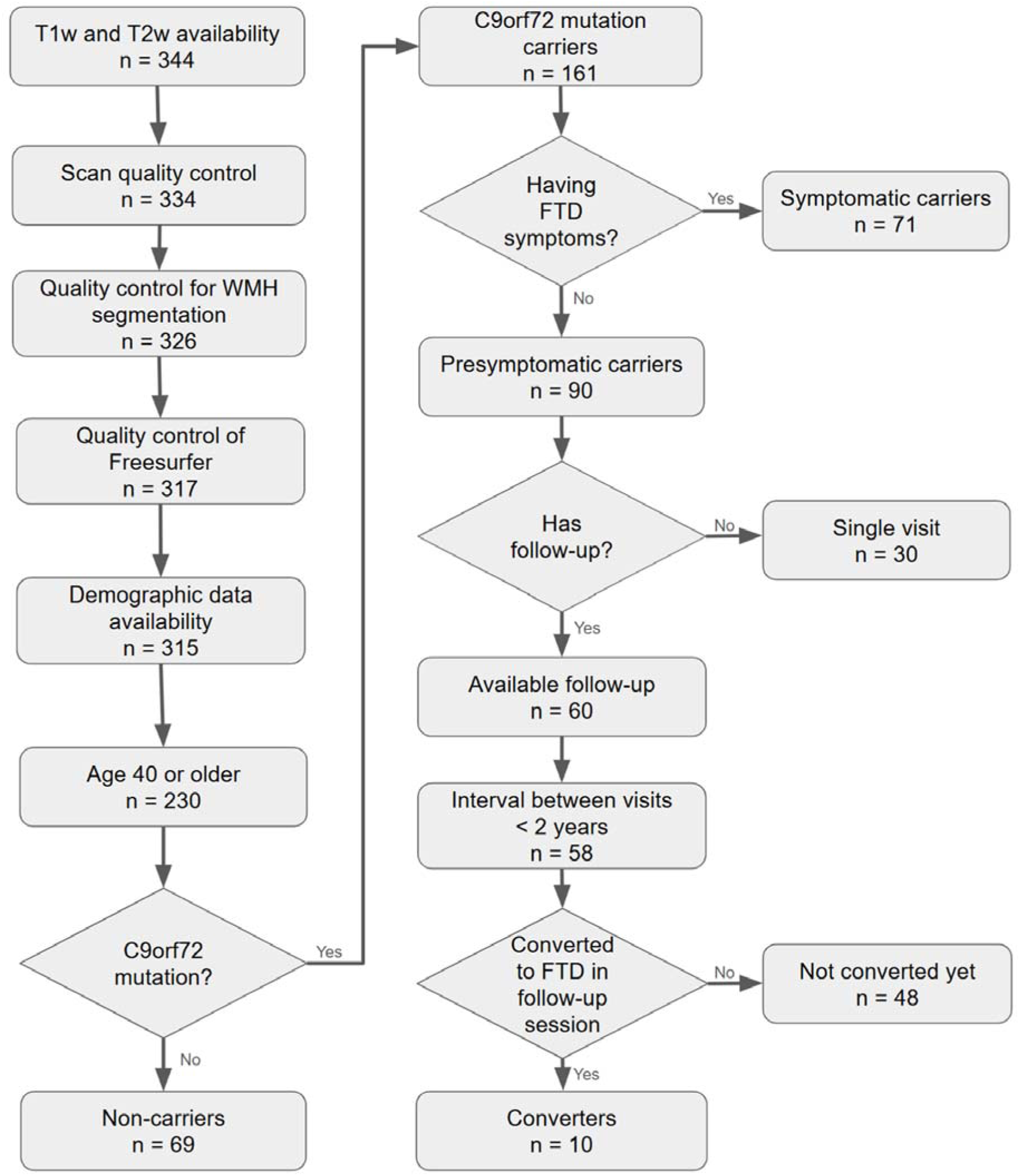
Flowchart of Inclusion and Exclusion Criteria. Illustrates the sequential application of criteria for participant selection, with the corresponding number of subjects retained at each step.

### 2.2 MRI Acquisition

T1-weighted and T2-weighted MRI scans were obtained on 3T scanners (Siemens Trio/Skyra/Prisma, Philips, and General Electric) across multiple GENFI sites. All sites followed harmonized acquisition protocols established by the GENFI study to ensure inter-site consistency and high image quality.^6^ T1-weighted imaging was acquired using sagittal 3D MPRAGE or equivalent sequences, and T2-weighted imaging using sagittal 3D fast spin echo sequences. Typical acquisition parameters included inversion time of approximately 850 ms, repetition time of 2000 ms, echo time of 3 ms, flip angle of 8°, slice thickness of 1.1 mm, and 200–208 slices per scan.

### 2.3 MRI Processing

#### 2.3.1 Cortical Thickness

T1-weighted images were processed using FreeSurfer v7.1.1 for cortical surface reconstruction and cortical thickness estimation. Parcellation was performed with the Desikan–Killiany atlas, yielding 34 regions per hemisphere. Mean cortical thickness was extracted for each region. All cortical and subcortical segmentations were visually inspected for accuracy, and subjects with inaccurate segmentations were excluded from further analysis (see Figure 1).

#### 2.3.2 Subcortical Segmentation

Subcortical structures were segmented using FreeSurfer’s ASEG pipeline, which applies a probabilistic atlas and Bayesian labeling framework. Extracted volumes included the thalamus, hippocampus, amygdala, caudate, putamen, pallidum, ventral diencephalon, lateral ventricles, third and fourth ventricles, brainstem, cerebellar cortex, cerebellar white matter, and corpus callosum subdivisions (anterior, central, mid-anterior, mid-posterior, posterior). These volumetric measures were included as features in the analysis.

#### 2.3.3 White Matter Hyperintensity Quantification

White matter hyperintensities (WMH) were segmented with BISON^21^ using combined T1- and T2-weighted data. Preprocessing included denoising, intensity normalization, and inter-modal registration; segmentation used a trained random forest, followed by lobar parcellation with the Hammers atlas. Visual quality control was performed, and outliers were excluded. Full methodological details are available in our prior work.^18^

### 2.4 Data Standardization and Feature Selection

All neuroimaging biomarkers, along with age and education, were standardized using z-scores, representing each value as the number of standard deviations from the mean. This approach facilitates comparability across different types of measurements with varying units and scales. The model incorporated a diverse set of features, including WMH volumes, cortical thickness measurements, brain segmentation values, and demographic variables (age, sex, handedness, education, employment status, and ethnicity). To account for inter-subject variability in head size, total intracranial volume (TIV) was also included as a covariate in our analysis.

Given the relatively small sample size, including all features in the model would risk overfitting. Therefore, dimensionality reduction was necessary to optimize model performance and enhance generalizability. Previous studies have demonstrated that applying feature selection or extraction prior to classification improves predictive accuracy,^22^ further motivating this step. To reduce dimensionality and identify the most informative components, principal component analysis (PCA) was applied. PCA transforms the original correlated variables into a smaller set of uncorrelated principal components, capturing the majority of the variance within the data.

### 2.5 Classifier Development and Validation

A random forest classifier was developed to distinguish symptomatic *C9orf72* mutation carriers from non-carrier controls based on neuroimaging and demographic data. The random forest algorithm is an ensemble learning method that constructs multiple decision trees during training and outputs the majority vote (i.e., the most frequent class) for classification tasks.^23^ This method was selected because it is inherently robust to overfitting, particularly when compared to individual decision trees, and performs well in settings where the number of variables (data dimensionality) exceeds the number of observations.^22,24^ Additionally, random forests are relatively interpretable and can handle complex interactions between features without strong parametric assumptions.^23,25^

To optimize the classifier’s performance, a grid search was performed to tune key hyperparameters, specifically focusing on the maximum tree depth and the number of estimators. After extensive evaluation, we found that a configuration with a maximum depth of one and 20 estimators provided the best performance. In this setup, the random forest consisted of 20 shallow decision trees, each limited to a single split. This simple structure was chosen to minimize overfitting, promote model interpretability, and maintain a high level of classification accuracy.

Model validation was conducted using stratified five-fold cross-validation. In this method, the dataset was divided into five equally sized folds, with each fold maintaining the same proportion of symptomatic and control samples as the full dataset. In each iteration, four folds were used for training and the remaining fold for testing. Stratification ensured that class imbalance did not bias the training or evaluation process. To further enhance the robustness of the results, the entire cross-validation procedure was repeated 100 times with random shuffling of the data before each repetition. This repeated cross-validation minimizes the impact of chance splits and improves the reliability of performance estimates.

The model was trained on the first 10 principal components obtained from the PCA. For each iteration, PCA was performed exclusively on the training folds, and the resulting transformation was then applied to the corresponding test fold. This approach ensured that the test data remained completely independent from the training process, avoiding any data leakage. Classification performance was assessed by calculating accuracy, sensitivity, and specificity across all iterations.

Figure 2 provides an overview of the study workflow. All machine learning analyses were conducted using the scikit-learn library (version 1.4.0) in Python (version 3.9.5).

**Figure 2.**
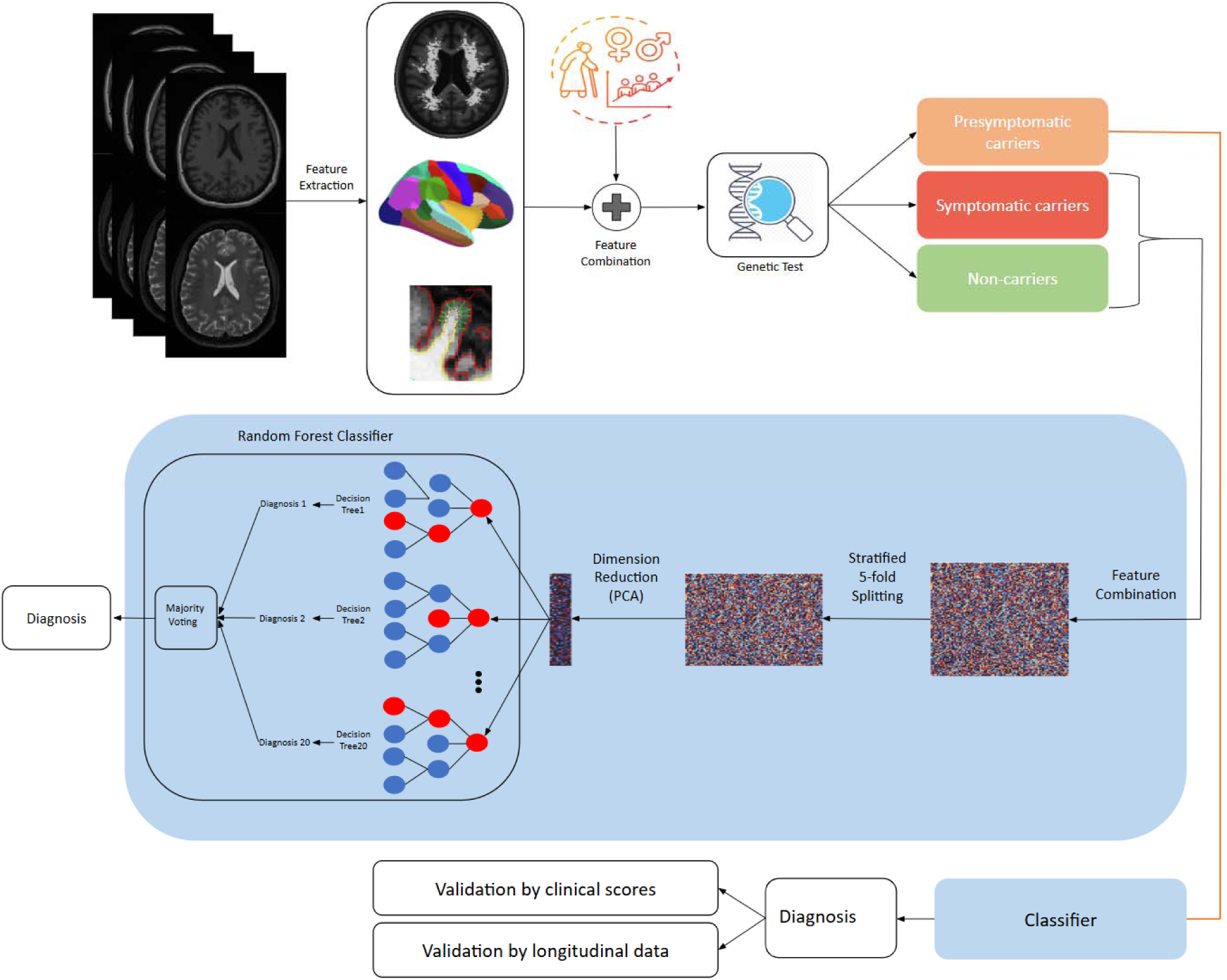
Overview of the study workflow. MRI data were preprocessed and used to derive imaging features. Dimensionality reduction was performed using PCA, and a random forest classifier was trained to distinguish symptomatic *C9orf72* mutation carriers from non-carrier controls with 5-fold cross-validation. The trained model was then applied to the presymptomatic cohort, and diagnostic predictions were validated using clinical scores and longitudinal follow-up data.

### 2.6 Model Interpretation with SHAP Analysis

To interpret the contribution of each neuroimaging and demographic feature to the classification, we employed SHapley Additive exPlanations (SHAP).^26,27^ SHAP values quantify the marginal contribution of each feature to the model’s prediction, enabling a global ranking of feature importance.

Because the classifier was trained on principal components rather than the raw features, we implemented SHAP analysis on the full pipeline using a wrapper function to map predictions back to the original feature space. A KernelExplainer was applied with the training data from each fold as the background distribution, and SHAP values were computed for all held-out test samples. This ensured that interpretation was based on completely unseen data and avoided data leakage.

For each cross-validation fold, SHAP values were calculated and then aggregated across all repetitions of the cross-validation procedure. From these, we derived mean absolute SHAP values, providing a ranked measure of each feature’s contribution to model predictions and enabling identification of the neuroimaging biomarkers most strongly associated with disease classification.

### 2.7 Application of the Classifier to Presymptomatic Carriers

#### 2.7.1 Classifier-Derived Risk Scoring

Following model training and validation (distinguishing symptomatic *C9orf72* carriers from healthy controls), the trained random forest classifier was applied to an independent cohort of presymptomatic *C9orf72* mutation carriers to evaluate its potential for identifying individuals at higher risk of imminent symptom onset. Each participant received a probabilistic disease score ranging from zero to one, where a score of zero indicated high similarity to healthy controls and a score of one indicated high similarity to symptomatic FTD patients. These scores were derived by averaging the binary output of our classifier across 100 iterations performed during the cross-validation phase.

Presymptomatic carriers with a mean disease score above 0.5 were categorized as high-risk, suggesting a greater likelihood of near-term conversion to symptomatic status. Those with a score below 0.5 were classified as low-risk. Because the classifier was trained exclusively on symptomatic carriers and non-carrier controls, its application to presymptomatic individuals served as an independent test, with no risk of data leakage.

#### 2.7.2 Neuropsychological and Biomarker Analyses

Because exact conversion times were unavailable for most presymptomatic participants, the clinical validity of the classifier was evaluated cross-sectionally at baseline. Neuropsychological performance and peripheral biomarkers were compared between high-risk and low-risk presymptomatic carriers (defined in Section 2.7.1). The neuropsychological battery^28^ included the following standardized measures: the Mini-Mental State Examination (MMSE; global cognition), Trail Making Test-Part B (TMT-B; executive function, task-switching, and cognitive flexibility), Digit Symbol Substitution Test (processing speed, attention, and working memory), Boston Naming Test (confrontation naming, word retrieval, and language), Verbal Fluency tasks (phonemic and semantic fluency), and the MiniSEA (Mini Social and Emotional Assessment; social cognition and emotional processing).

Neurofilament light chain (NfL), a peripheral blood marker of axonal degeneration, was measured to validate classifier predictions through group comparisons and correlations with disease scores.

In addition to group comparisons, associations between the continuous baseline disease score and each neuropsychological measure, and biomarker level were assessed using correlation analyses. All analyses were conducted in the independent presymptomatic cohort used for external application of the classifier. All analyses were conducted in the independent presymptomatic cohort used for external application of the classifier.

#### 2.7.3 Longitudinal Validation

To further assess the true predictive validity of the classifier for imminent clinical conversion, longitudinal data were analyzed from 58 presymptomatic *C9orf72* carriers with at least one follow-up visit within 24 months of baseline. Among these, 10 individuals converted to symptomatic status during follow-up. Conversion was defined as the onset of clinical features consistent with an FTD-spectrum disorder by expert clinical consensus.

As mentioned in Section 2.7.1, our model categorized participants into low-risk and high-risk groups. Predictive validity was evaluated by comparing classifier-assigned risk status against observed conversion outcomes: converters classified as high-risk were considered true positives, converters classified as low-risk as false negatives, non-converters classified as high-risk as false positives, and non-converters classified as low-risk as true negatives. This framework enabled estimation of sensitivity, specificity, and positive and negative predictive values of the classifier for predicting conversion within two years.

## 3 Results

### 3.1 Participant Overview

Our sample included 69 non-carrier controls, 71 symptomatic carriers, and 90 presymptomatic carriers of the *C9orf72* mutation. Demographic characteristics are summarized in Table 1. The symptomatic group was significantly older than both presymptomatic carriers and controls, and the proportion of male participants was higher in the symptomatic group. Presymptomatic carriers had higher education levels than symptomatic carriers. Accordingly, age, sex, and education were treated as confounding factors in the diagnostic model. In terms of sample size, the symptomatic and control groups were comparable, providing a balanced training dataset. In the symptomatic cohort, 70.4% were diagnosed with bvFTD, 19.7% with ALS or FTD-ALS, 5.6% with PPA, and 4.2% with other dementia syndromes.

**Table 1.** Demographic characteristics of the study cohort.

| Cohort | Healthy controls | Presymptomatic carriers | Symptomatic carriers |
| --- | --- | --- | --- |
| Number of subjects | 69 | 90 | 71 |
| Sex, male (%) | 28 (40.58%) | 34 (37.77%) | 43 (60.56%) |
| Age (mean ± std) | 52.92 ± 9.54 | 51.18 ± 8.05 | 64.68 ± 7.48 <sup>a</sup> |
| Education (mean ± std) | 13.86 ± 2.88 | 14.21 ± 3.16 | 12.89 ± 3.56 <sup>b</sup> |

Values are reported as mean ± standard deviation or number (percentage).
*a*: Symptomatic carriers were significantly older than both presymptomatic carriers and healthy controls (Mann–Whitney U = 5621, Bonferroni-corrected $p < 0.001$ ).
*b*: Presymptomatic carriers had significantly higher education levels than symptomatic carriers (Mann–Whitney U = 2389.5, Bonferroni-corrected $p = 0.017$ ).

### 3.2 Model and Classifier Performance

We retained the first 10 principal components from the PCA, as we found that n=10 provides the best compromise between keeping the number of components small and explaining a good portion of the total variance (75%). The random forest classifier demonstrated strong performance in distinguishing symptomatic *C9orf72* mutation carriers from non-carrier controls. Averaged across 100 cross-validation iterations, the model achieved an accuracy of 87.0% ± 2.52%, a sensitivity of 83.82% ± 3.98% (correctly identifying symptomatic carriers), and a specificity of 90.38% ± 3.91% (correctly identifying controls). The corresponding ROC curve is presented in Figure 3, indicating robust discriminative ability.

**Figure 3.**
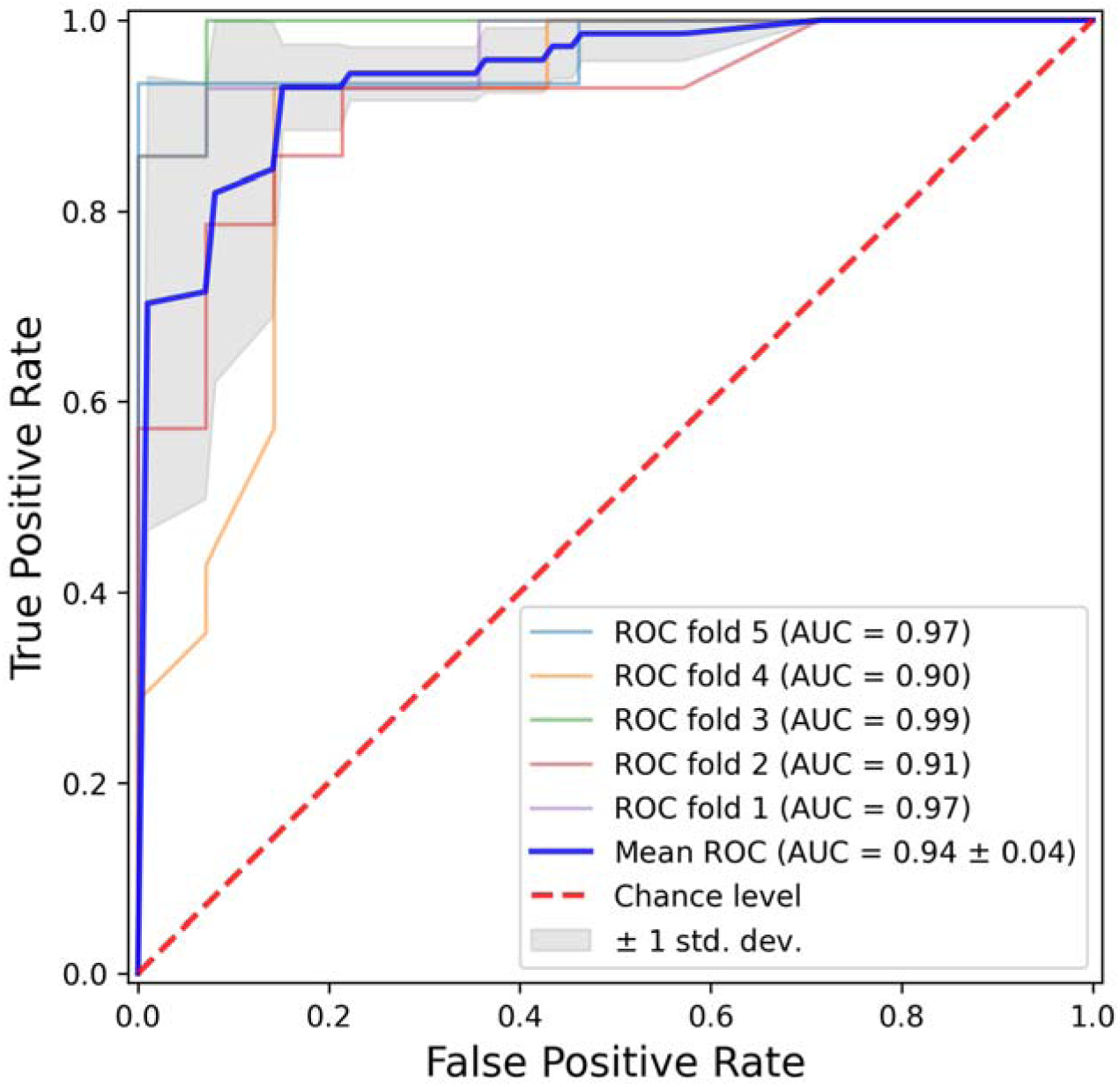
Receiver operating characteristic (ROC) curves for classifier performance. ROC curves illustrate the model’s ability to distinguish symptomatic C9orf72 carriers from healthy controls. Curves are shown for each fold of the 5-fold cross-validation, along with the mean ROC curve and ±1 standard deviation shaded region. The red dashed line represents chance-level classification.

### 3.3 Application to Presymptomatic Carriers

Applying the trained classifier to the presymptomatic cohort, 19 out of 90 individuals (21.1%) were categorized as high-risk based on their neuroanatomical similarity to symptomatic FTD cases, as measured by our classifier. The remaining 71 individuals (78.9%) were classified as low-risk, reflecting greater neuroanatomical resemblance to healthy controls (with age, sex, and education already included as model features, thereby controlling for these demographic factors).

All reported p-values (p_FDR_) are adjusted for multiple comparisons using the Benjamini– Hochberg FDR procedure.

#### 3.3.1 Neuropsychological and Biomarker Differences Between Risk Groups

As summarized in Table 2 and Figure 4, neuropsychological testing revealed significantly worse cognitive performance among high-risk individuals across several domains related to FTD:

- Trail Making Test B: High-risk individuals showed significantly slower performance (p_FDR_ = 0.005), indicating mild executive weaknesses.
- Verbal Fluency Task: Lower word production in the high-risk group (p_FDR_ = 0.016), suggesting reduced lexical access and semantic memory.
- Digit Symbol Substitution Test: Markedly reduced scores in high-risk individuals (p_FDR_ = 0.002), reflecting lower processing speed and attention.
- MiniSEA: Significantly lower scores in the high-risk group (p_FDR_ = 0.016), consistent with early weaknesses in social and emotional cognition.
- Boston Naming Test: High-risk participants showed a trend toward lower naming accuracy, suggesting emerging language difficulties; however, this effect did not remain significant after FDR correction (p_FDR_ = 0.064).

**Figure 4.**
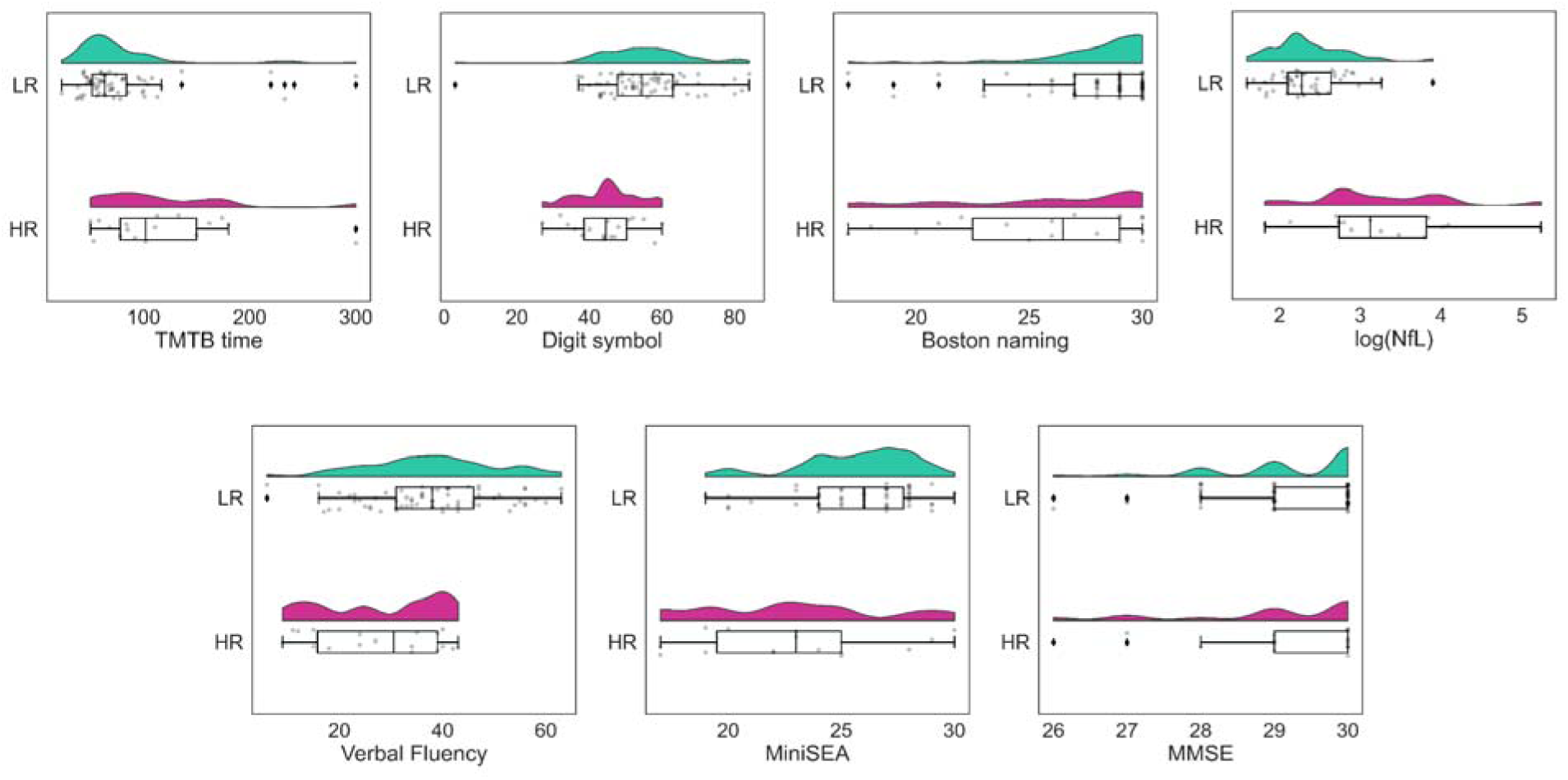
Clinical and biomarker differences between high- and low-risk presymptomatic *C9orf72* carriers. Comparison between high-risk (HR) and low-risk (LR) groups of presymptomatic *C9orf72* mutation carriers, categorized based on model-derived predictions. The figure illustrates group differences in clinical performance across neuropsychological tests, as well as levels of neurofilament light. LR: Low-Risk; HR: High-Risk.

**Table 2.**
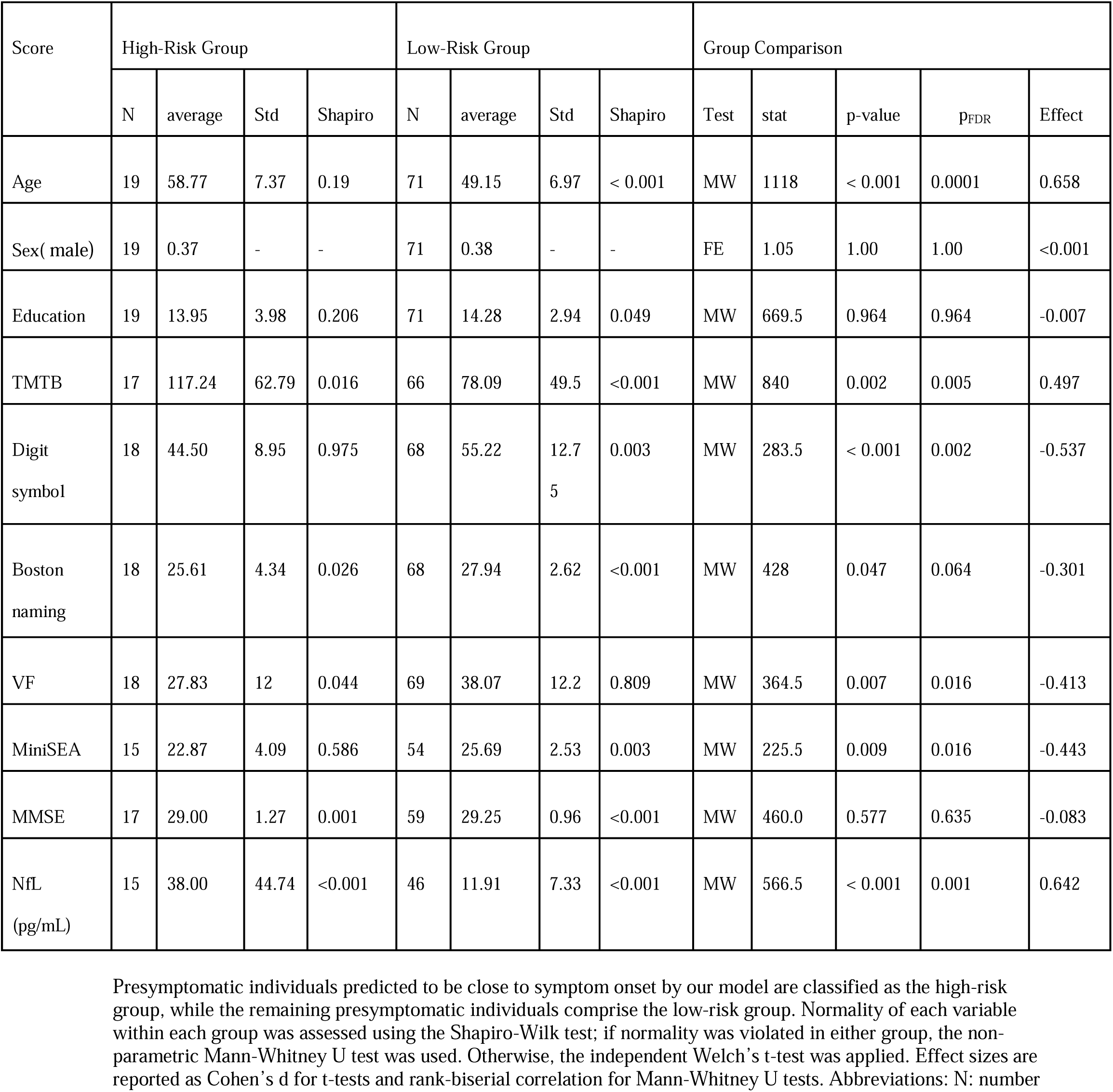
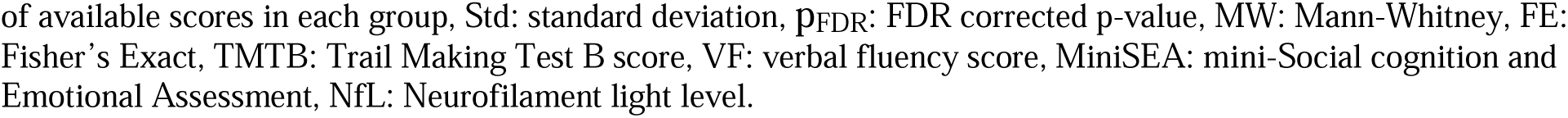
Comparison of neuropsychological scores and blood biomarker between high- and low-risk presymptomatic carriers.

| Score | High-Risk Group |  |  |  | Low-Risk Group |  |  |  | Group Comparison |  |  |  |  |
| --- | --- | --- | --- | --- | --- | --- | --- | --- | --- | --- | --- | --- | --- |
|  | N | average | Std | Shapiro | N | average | Std | Shapiro | Test | stat | p-value | p<sub>FDR</sub> | Effect |
| Age | 19 | 58.77 | 7.37 | 0.19 | 71 | 49.15 | 6.97 | < 0.001 | MW | 1118 | < 0.001 | 0.0001 | 0.658 |
| Sex( male) | 19 | 0.37 | - | - | 71 | 0.38 | - | - | FE | 1.05 | 1.00 | 1.00 | <0.001 |
| Education | 19 | 13.95 | 3.98 | 0.206 | 71 | 14.28 | 2.94 | 0.049 | MW | 669.5 | 0.964 | 0.964 | -0.007 |
| TMTB | 17 | 117.24 | 62.79 | 0.016 | 66 | 78.09 | 49.5 | <0.001 | MW | 840 | 0.002 | 0.005 | 0.497 |
| Digit symbol | 18 | 44.50 | 8.95 | 0.975 | 68 | 55.22 | 12.75 | 0.003 | MW | 283.5 | < 0.001 | 0.002 | -0.537 |
| Boston naming | 18 | 25.61 | 4.34 | 0.026 | 68 | 27.94 | 2.62 | <0.001 | MW | 428 | 0.047 | 0.064 | -0.301 |
| VF | 18 | 27.83 | 12 | 0.044 | 69 | 38.07 | 12.2 | 0.809 | MW | 364.5 | 0.007 | 0.016 | -0.413 |
| MiniSEA | 15 | 22.87 | 4.09 | 0.586 | 54 | 25.69 | 2.53 | 0.003 | MW | 225.5 | 0.009 | 0.016 | -0.443 |
| MMSE | 17 | 29.00 | 1.27 | 0.001 | 59 | 29.25 | 0.96 | <0.001 | MW | 460.0 | 0.577 | 0.635 | -0.083 |
| NfL (pg/mL) | 15 | 38.00 | 44.74 | <0.001 | 46 | 11.91 | 7.33 | <0.001 | MW | 566.5 | < 0.001 | 0.001 | 0.642 |

Presymptomatic individuals predicted to be close to symptom onset by our model are classified as the high-risk group, while the remaining presymptomatic individuals comprise the low-risk group. Normality of each variable within each group was assessed using the Shapiro-Wilk test; if normality was violated in either group, the non-parametric Mann-Whitney U test was used. Otherwise, the independent Welch's t-test was applied. Effect sizes are reported as Cohen's d for t-tests and rank-biserial correlation for Mann-Whitney U tests. Abbreviations: N: number
of available scores in each group, Std: standard deviation, $p_{\text{FDR}}$ : FDR corrected p-value, MW: Mann-Whitney, FE: Fisher's Exact, TMTB: Trail Making Test B score, VF: verbal fluency score, MiniSEA: mini-Social cognition and Emotional Assessment, NfL: Neurofilament light level.

There was no significant difference in MMSE scores, suggesting that global cognitive impairment had not yet emerged, in line with the presymptomatic status of all participants.

Consistent with these neuropsychological findings, NfL levels were significantly elevated in the high-risk group (p_FDR_ = 0.001), reflecting greater underlying neurodegeneration.

#### 3.3.2 Correlation between Disease-Score as a Continuous Measure and Clinical/Biomarker Measures

Pearson correlation analyses revealed significant associations between the classifier’s continuous disease scores and a range of clinical and biomarker measures. Higher disease scores correlated with worse cognitive performance and higher NfL levels, supporting the biological validity of the model’s predictions. Detailed correlation values are presented in Table 3.

**Table 3.** Correlation between model-derived disease score and clinical or biomarker measures.

| Score | correlation | P-value | FDR corrected P-value |
| --- | --- | --- | --- |
| TMTB | 0.36 | < 0.001 | 0.001 |
| Digit symbol | - 0.40 | < 0.001 | 0.001 |
| Boston naming | - 0.35 | 0.001 | 0.001 |
| VF | - 0.37 | < 0.001 | 0.001 |
| MiniSEA | - 0.39 | < 0.001 | 0.001 |
| NfL | 0.44 | < 0.001 | 0.001 |
| MMSE | - 0.09 | 0.432 | 0.432 |

Pearson correlation analysis between clinical/blood scores and the disease score estimated by our model for each presymptomatic mutation carrier. Abbreviations: TMTB: Trail Making Test B, Digit Symbol Substitution Test, Boston Naming Test, VF: Verbal Fluency, MiniSEA: Social and Emotional Assessment, NfL: Neurofilament Light, MMSE: Mini-Mental State Examination.

### 3.4 Longitudinal Validation and Prediction of Conversion

To assess the model’s real-life predictive utility, we focused on the 58 presymptomatic cases with clinical follow-up visits within two years of their baseline MRI assessment. 10 individuals (17.2%) converted to symptomatic FTD during the follow-up period, as determined by expert clinical consensus, while the remaining 48 individuals (82.8%) remained asymptomatic. Among converters, 30% were diagnosed with bvFTD, 30% with ALS or FTD-ALS, and 40% with other FTD-spectrum syndromes.

At baseline, the classifier identified 13 individuals (22.4%) as high-risk based on their neuroanatomical profiles. Among these, 7 subsequently converted to symptomatic status, resulting in a sensitivity of 70% and a positive predictive value of 53.8%. The remaining 45 individuals (77.6%) were classified as low-risk, of whom 42 remained asymptomatic, corresponding to a specificity of 87.5% and a negative predictive value of 93.3%. Of the 3 converters in the low-risk group, none had a typical bvFTD/PPA clinical profile (1 ALS and 2 other types of syndromes). Overall, the model achieved a prediction accuracy of 84.5% in distinguishing converters from non-converters over the two-year interval.

### 3.5 Model Interpretation and Feature Importance

Global feature importance, quantified as mean absolute SHAP value, revealed a multimodal profile spanning cortical thickness, subcortical volumes, cerebellar structures, ventricular size, and white matter hyperintensities (Figure 5). SHAP values quantify each feature’s contribution to the model’s prediction, with larger values indicating greater influence. The most influential feature was right paracentral cortical thickness, followed by left lateral orbitofrontal thickness and left cortical volume. Other high-ranking cortical measures included left postcentral, right rostral middle frontal, bilateral precentral, left medial orbitofrontal, left frontal pole, right superior parietal, and left caudal middle frontal thickness, indicating prominent involvement of the sensorimotor cortex (perirolandic strip) and prefrontal areas.

**Figure 5.**
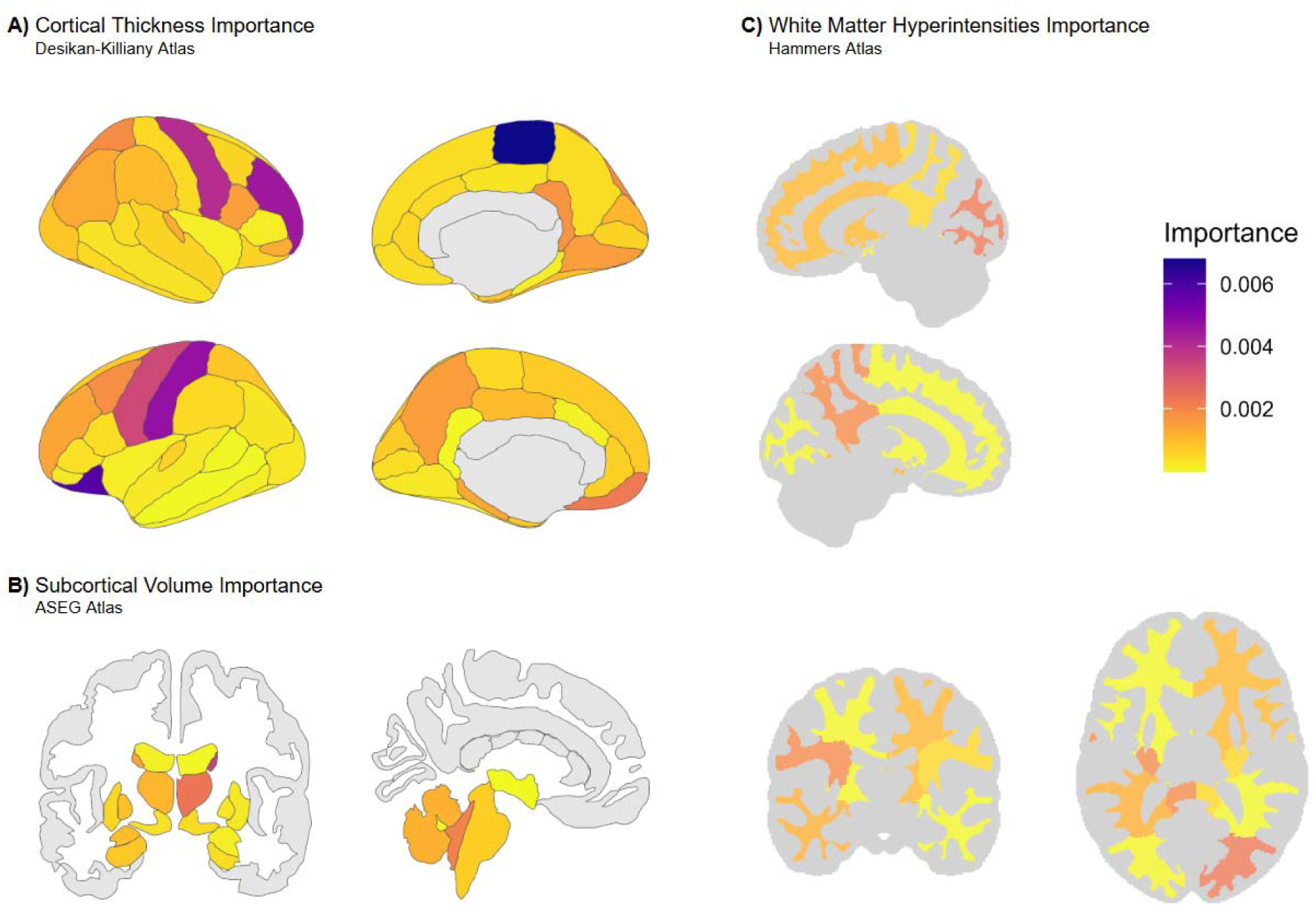
Feature importance derived from SHAP analysis. Mean absolute SHAP values are shown for different neuroimaging features contributing to model predictions. Larger values indicate features with greater influence on the model’s output. (A) Cortical thickness across regions defined by the DKT atlas (first row: right hemisphere; second row: left hemisphere). (B) Subcortical volumes based on the ASEG atlas. (C) White matter hyperintensities in different lobes (first row: right hemisphere; second row: left hemisphere). Mean absolute SHAP values ranged from ∼0.00001 to ∼0.0068.

Subcortical structures also featured prominently among the top 20, including the right caudate, right thalamus, and left cerebellar white matter volumes. In addition, the fourth ventricle volume contributed to the overall feature profile.

WMH were highly informative in the model. Both total WMH volume and lobar measures (right occipital WMH and left parietal WMH) ranked within the top 20, indicating that white matter signal changes carried substantial discriminative value alongside cortical and subcortical markers.

## 4 Discussion

Predicting the age at symptom onset in genetic FTD is a critical challenge, yet one with profound implications for personalized medicine and clinical trial design. Prior evidence demonstrates that neurodegenerative changes, particularly structural brain alterations, can precede overt clinical symptoms by years. These subclinical changes offer a unique opportunity to identify biomarkers that signal proximity to phenoconversion, especially in genetically at-risk individuals. However, the complex and heterogeneous nature of FTD pathology suggests that relying on a single neuroimaging biomarker may be insufficient for accurate prognosis. In this study, we addressed this challenge by leveraging a multi-modal machine learning approach to integrate multiple neuroanatomical features and derive a comprehensive “brain fingerprint” of *C9orf72*-associated FTD.

Our MRI-based machine learning classifier demonstrated high classification accuracy to distinguish symptomatic FTD cases from controls, indicating that a robust neuroanatomical signature of FTD can be extracted using dimensionality-reduction and ensemble learning techniques despite the heterogeneity of clinical syndromes. When applied to a separate group of presymptomatic carriers, the high-risk presymptomatic individuals identified by the classifier showed significantly poorer performance on several key neuropsychological measures related to FTD, including executive function, language, and social cognition, despite being considered clinically asymptomatic by experts. This finding supports the validity of our model in identifying individuals on the cusp of phenoconversion. Furthermore, longitudinal follow-up confirmed the model’s prognostic utility, with an accuracy of 84.5% in predicting conversion to symptomatic FTD within two years. Individuals predicted as negative by the classifier were very unlikely to convert to symptomatic FTD within two years, highlighting the model’s strong negative predictive value.

Our SHAP analysis provided a data-driven ranking of the most informative features, revealing a complex neuroanatomical signature. Both cortical and subcortical measures, along with WMH, played a critical role in distinguishing *C9orf72*-FTD from controls. Cortical thickness in the paracentral lobule, precentral/postcentral gyri, and orbitofrontal regions emerged as particularly important. This is consistent with a growing body of literature highlighting peri-rolandic (primary motor) and frontal involvement in *C9orf72*-FTD.^9,10,16,29^ The prominent role of the paracentral lobule underscores potential subclinical motor-system involvement even in the absence of a co-morbid ALS diagnosis, while contributions from orbitofrontal and rostral middle frontal areas align with the behavioral variant FTD phenotype.^7,30^

Subcortical contributions, particularly from the thalamus and caudate, reinforce evidence that thalamo-striatal circuits are a key site of pathology in *C9orf72*-FTD.^10–12^ Cerebellar and fourth-ventricle measures also contributed to classification, consistent with reports of cerebellar and ventricular changes in *C9orf72*-FTD.^6,19,31^

Importantly, WMH measures were among the most informative features in our model. This finding aligns with recent evidence that *C9orf72* carriers demonstrate elevated WMH burden, including during presymptomatic phases.^18^ This suggests that WMH may represent an underappreciated imaging biomarker in the disease, complementing gray matter and subcortical measures. Overall, the complexity of neuroanatomical features identified in this study supports our multi-feature machine learning approach as providing a more holistic view of the disease process compared to single-marker studies.

This study was designed to evaluate the potential of a neuroimaging-informed machine learning approach for risk stratification. We acknowledge that the specific model and hyperparameters we selected, while effective, may not be the optimal combination. Exploring the vast space of alternative machine learning models, feature engineering techniques, and hyperparameter tunings was beyond the scope of this study. Our main objective was to demonstrate that a machine learning approach could yield a promising performance in identifying high-risk converters with a single diagnostic tool of MRI, which our results strongly support. Future work could systematically explore other model architectures and tuning strategies to potentially enhance performance. Our analysis was also limited to a specific set of structural neuroimaging features. Future studies could integrate other imaging modalities, such as diffusion tensor imaging (DTI) or functional MRI (fMRI), which may capture complementary aspects of disease pathology, but tend to be harder to harmonize at the individual level. We acknowledge the age difference between our symptomatic FTD and control cohorts. Due to the relatively small sample size, age-matching or subsampling was not feasible. We addressed this by including age as a feature in our classifier. Importantly, our SHAP analysis indicated that age did not play a significant role in differentiating FTD from the healthy control group, suggesting that this age difference did not introduce a major bias into our primary classification results.

There is also an inherent limitation to the establishment of a precise time for phenoconversion. First, the onset of symptoms in FTD is often subtle and gradual, complicating efforts to anchor disease progression to a specific time point. Second, the exact timing of diagnosis is dependent on the frequency of clinical visits, and initial symptoms may be misattributed or missed entirely, leading to inconsistencies in diagnostic timing across different clinical sites and participants.

While the size of the longitudinal cohort was limited, our use of detailed neuropsychological assessments allowed us to detect meaningful subclinical differences between high- and low-risk individuals. The inclusion of participants from multiple international sites strengthens the generalizability and robustness of our findings, demonstrating the possibility for this approach to be applied in a multi-center context.

While our findings highlight the value of neuroimaging-based prediction, the literature has also emphasized the importance of other biomarkers in the diagnosis of FTD. One of the most studied is NfL, which rises sharply around the time of symptom onset,^32^ and has shown good performance in detecting converters, though reported sensitivity is low. For instance, Linnemann et al.^33^ observed 43% sensitivity and 100% specificity in a GENFI cohort. Of note, their study merged all mutations together, and accuracy could be lower in *C9orf72* carriers, where NfL rises slowly over a longer period presymptomatically.^34^ In contrast, our neuroimaging based method achieved higher sensitivity, making it valuable for identifying individuals earlier in the disease course. Future studies should therefore explore combining imaging and fluid biomarkers to improve predictive power.

Finally, our focus on the *C9orf72* mutation was motivated by its distinct disease trajectory, characterized by early and widespread neuroanatomical changes,^6,10^ making it an ideal model for identifying early imaging biomarkers. This approach may serve as a template for future studies targeting other genetic variants, such as *MAPT* and *GRN*, where early detection is equally critical.

In summary, our findings highlight the potential of neuroimaging-informed machine learning to identify individuals at high risk of conversion to FTD within a two-year period. This represents an important step toward transforming MRI, an accessible and non-invasive tool, into a predictive biomarker of disease onset. Beyond its diagnostic value, this approach could inform the design of future clinical trials by enabling the selective enrollment of asymptomatic mutation carriers identified as high-risk based on their brain signature. Clinically, it may also provide a useful framework for delivering individualized prognostic information to patients and their families.

## Data availability

The data used in this study is part of the GENFI dataset and can be accessed upon reasonable request through the study website (www.genfi.org), subject to review and approval by the GENFI data access committee.

## Acknowledgements

We would like to thank all participants and their families for taking part in the GENFI study.

## Appendix 1

### GENFI consortium members

Annabel Nelson, Martina Bocchetta, David Cash, David L Thomas, Emily Todd, Hanya Benotmane, Jennifer Nicholas, Kiran Samra, Rachelle Shafei, Carolyn Timberlake, Thomas Cope, Timothy Rittman, Antonella Alberici, Enrico Premi, Roberto Gasparotti, Valentina Cantoni, Emanuele Buratti, Andrea Arighi, Chiara Fenoglio, Elio Scarpini, Giorgio Fumagalli, Vittoria Borracci, Giacomina Rossi, Giorgio Giaccone, Giuseppe Di Fede, Paola Caroppo, Pietro Tiraboschi, Sara Prioni, Veronica Redaelli, David Tang-Wai, Ekaterina Rogaeva, Miguel Castelo-Branco, Morris Freedman, Ron Keren, Sandra Black, Sara Mitchell, Christen Shoesmith, Robart Bartha, Rosa Rademakers, Janne M. Papma, Lucia Giannini, Rick van Minkelen, Yolande Pijnenburg, Benedetta Nacmias, Camilla Ferrari, Cristina Polito, Gemma Lombardi, Valentina Bessi, Michele Veldsman, Christin Andersson, Hakan Thonberg, Linn Öijerstedt, Vesna Jelic, Paul Thompson, Tobias Langheinrich, Albert Lladó, Anna Antonell, Jaume Olives, Mircea Balasa, Nuria Bargalló, Sergi Borrego-Ecija, Ana Verdelho, Ana Gorostidi, Jorge Villanua, Marta Cañada, Mikel Tainta, Miren Zulaica, Myriam Barandiaran, Patricia Alves, Benjamin Bender, Lisa Graf, Annick Vogels, Mathieu Vandenbulcke, Philip Van Damme, Rose Bruffaerts, Koen Poesen, Pedro Rosa-Neto, Serge Gauthier, Anne Bertrand, Aurélie Funkiewiez, Daisy Rinaldi, Dario Saracino, Olivier Colliot, Sabrina Sayah, Catharina Prix, Elisabeth Wlasich, Olivia Wagemann, Sandra Loosli, Sonja Schönecker, Tobias Hoegen, Jolina Lombardi, Sarah Anderl-Straub, Adeline Rollin, Gregory Kuchcinski, Maxime Bertoux, Thibaud Lebouvier, Vincent Deramecour, Beatriz Santiago, Diana Duro, Maria João Leitão, Maria Rosario Almeida, Miguel Tábuas-Pereira, Sónia Afonso

